# Analysing drivers of worldwide tidal wetland change

**DOI:** 10.1101/2024.08.27.609936

**Authors:** Lucie Perrodin, Alejandro Navarro, Maren Toor, Robert Canto, Madison Becker, Yanzhu Dong, Thomas Worthington, Nicholas J. Murray

## Abstract

Tidal wetlands are dynamic coastal ecosystems that can change in extent in response to a broad range of change drivers. We use high spatial resolution satellite imagery to estimate the relative influence of 18 classes of change drivers on observed tidal wetland gains and losses from 1999 to 2019, differentiating direct drivers as those observable at the site of ecosystem change, and indirect drivers as broader processes that influence changes without being directly visible. We developed a random sample of 2823 change detections from a global dataset of tidal wetland change and allocated each change event to driver classes using high-resolution time-series imagery. We identified that indirect drivers were the most widespread type of driver of tidal wetland change (70.9%), with flooding being the predominant driver for losses (47.5%) and unknown natural processes of change for gains (62.7%). Drivers often associated with climate change were evident in interpretations of wetland drivers, with increases in flooded area and reductions in vegetation cover suggesting the effects of relative sea level rise on tidal wetlands are observable in many areas. Our temporal analysis revealed that over 20 years, indirect drivers consistently contributed to larger proportions of gains and losses compared to direct drivers. Asia was the only continent where direct drivers of loss, such as agriculture (22.9%) and aquaculture (17.1%), outweighed indirect drivers, providing further evidence of the widespread transformation of Asia’s natural coastal ecosystems to anthropogenic shorelines. Globally, coastal land reclamations were mostly observed in mangrove ecosystems, where more than half of the observed losses were of anthropogenic origin. The most observed direct drivers of gains were altered land management and restoration, but none of them contributed to more than 5% of the total gains over 20 years. Our findings suggest a need for efficient conservation measures that allow the dynamic processes that characterise coastal ecosystems to persist, while simultaneously reducing the worldwide impact of direct human activities.

## 1. Introduction

Losses of coastal ecosystems, including tidal flat, mangrove and tidal marsh ecosystems (hereafter collectively termed ‘tidal wetlands’), have been occurring for centuries. Recent global estimates suggest that around 30% of mangroves and 16% of tidal flats have been lost over the last 40 years, threatening the delivery of essential ecosystem services (1984-2016; Onrizal et al, 2018; Murray et al, 2019). The global remote sensing analyses that enabled these estimates have revealed the important role that wetland gains also play in global net change, including an estimated 1921 km^2^ of tidal wetland gain over the same period (Murray et al, 2022).

However, there remains a lack of comprehensive knowledge regarding the complex processes that are associated with the losses and gains of tidal wetlands that have been detected worldwide. Recent studies into the dynamics of global tidal wetland extent often use remote sensing classifications of detected change that cannot distinguish the complex and often subtle drivers of that change. Furthermore, these studies suffer from simplistic classification schemes which fail to account for the full diversity of known drivers of coastal ecosystem change. A better understanding of the myriad drivers of both tidal wetland gains and losses is crucial to improve forecasting of environmental impacts and develop effective conservation strategies worldwide.

Globally, coastal development is widely recognised as the primary anthropogenic driver of tidal wetland losses (Cui et al, 2016; Newton et al, 2020). Coastal ‘reclamation’, the term generally applied to the process of converting areas of tidal wetlands to alternative land uses, has been responsible for an estimated 62% of global mangrove losses recorded from 2000 to 2016 (Thomas and al., 2017; Golberg et al, 2020; Turschwell et al. 2021). Murray et al (2022) suggested that 39% of detected tidal losses worldwide were the result of direct human activities (Murray et al, 2022).

In addition to drivers of wetland losses, there are a variety of change drivers that can result in gains of tidal wetland extent. Restoration and afforestation actions, such as mangrove planting, have enabled considerable gains in tidal wetlands in some areas (Bayraktarov et al, 2016; Lee et al, 2019). Actions such as the construction of dikes are also frequently used to trap sediments to maintain tidal wetland extent, but often ultimately lead to gains (Zhi et al, 2023). These tidal wetland management and restoration actions were estimated to account for nearly 14% of tidal wetland gains worldwide over the last two decades (Murray et al, 2022).

Owing to complex biophysical feedback mechanisms and natural coastal processes, tidal wetlands can, unlike many other ecosystem types, naturally gain or lose extent in response to a broad range of natural processes (Ciro Aucelli et al, 2017; Goldberg et al, 2020). For example, the physical processes of erosion and subsidence, as well as the natural transport of coastal sediments, can quickly result in losses in one area that are rapidly offset by gains in other areas. Sediment trapping by coastal vegetation can similarly enable tidal wetlands to maintain extent in the face of rising sea levels and migrate to higher elevations where conditions allow (Kirwan and Megonigal 2013). These processes that cause loss or gain often occur directly at the site of loss or gain (e.g. erosion) or are influenced by remote processes (e.g. sea level rise).

We focus this study on quantitatively characterising these changes to better understand the proximate and remote drivers of tidal wetland change. We assess the drivers of gains and losses of tidal wetlands between 1999 and 2019 based on high resolution image interpretation of a random sample of tidal wetland losses and gains obtained from the Global Tidal Wetlands Change (GTWC) data (Murray et al. 2022). We conducted analyses with the aim of understanding the global distribution and relative occurrence of drivers within each tidal wetland ecosystem type. To investigate important temporal dynamics of tidal wetland change, we used the GTWC data to observe the evolution over time of each driver’s contribution to changes. Our detailed assessment of global tidal wetland change, incorporating a comprehensive driver classification and analysis of change pace, will inform and enhance coastal wetland conservation and management efforts worldwide amidst rapid coastal changes.

## 2. Methods

### 2.1 Driver Classification Scheme

We assessed the relative influence of a suite of direct and indirect drivers of observed changes on tidal wetland extent. We focused on estimating the impacts of specific human activities that directly affect tidal wetland extent, such as aquaculture, urban development and dredging, while accounting for a range of natural drivers of change that are also known to influence tidal wetland extents over time (Li et al, 2014). As a global analysis, high resolution satellite imagery provides the basis for analysing observed wetland changes according to the types of driver that caused them, with interpretations of land cover types before, during and after tidal wetland change events forming the basis of our study.

To develop a driver classification scheme that accounts for the full diversity of direct and indirect drivers known to operate in coastal ecosystems, we reviewed previous field and remote sensing studies of coastal ecosystem change. Our qualitative review included more than 190 peer-reviewed publications that formed the basis of the development of our driver classification scheme. This analysis yielded 18 tidal wetland change drivers that have been shown to lead to losses or gains of extent of mangroves, tidal flats or saltmarsh around the world (Table 1).

**Table 1:** Description of the categories of drivers associated with their type and their effect on the extent. Note: owing to the inability to directly observe indirect drivers of change, each indirect change sample was annotated with its response, which we defined as the observed land cover type following a tidal wetland change event.

| Category | Description | Driver type | Effect on extent |
| --- | --- | --- | --- |
| Drying | Tidal wetland area is reduced or lost due to the reduction or elimination of a water source, such as tidal access | Indirect | Loss |
| Flooding | The ecosystem is replaced by permanent water, can be either saltwater or freshwater | Indirect | Loss |
| Sediment deposition | Observable addition of sediments to an initially non-intertidal zone | Indirect | Gain |
| Terrestrial conversion | Transitions to non-tidal land, possibly with terrestrial vegetation, beach, or bare land | Indirect | Loss |
| Unknown natural process | Change resulting from remote natural processes, that create conditions for the ecosystem to expand or disappear (e.g., breaching of sand banks) or that could not be clearly identified (e.g., vegetation gain/loss) | Indirect | Gain or Loss |
| Agriculture development | Organised farming of any scale and any type, including plantations | Direct | Gain or Loss |
| Altered land management | Managed realignment, land conversion, abandoned land (evidence of former activity) | Direct | Gain |
| Aquaculture development | Geometrically shaped ponds for any type and scale of aquaculture | Direct | Gain or Loss |
| Channelling/runnelling | Development of canals or ditching (mosquitos) through tidal wetlands | Direct | Gain or Loss |
| Dredging | Direct removal of tidal wetland or adjacent dredging that alters hydrology leading to erosion | Direct | Loss |
| General development | Bare land, cleared land, unidentifiable or unfinished development | Direct | Gain or Loss |
| Industrial development | Infrastructure (e.g, buildings or warehouses) built for industrial purposes | Direct | Loss |
| Mowing | Removal of mangrove or saltmarsh vegetation for personal, aesthetic, or management purposes | Direct | Loss |
| Ports/marinas | Platform/infrastructure capable of accommodating boats and ships | Direct | Gain or Loss |
| Restoration | Vegetation creation through construction and backfilling of soil, coastal engineering to promote vegetation growth, visible planting of vegetation, hydrologic restoration | Direct | Gain |
| Seawalls/dikes/levees | Seawalls, dikes, levees which do not belong to another infrastructure | Direct | Gain or Loss |
| Transport infrastructure | Terrestrial transport infrastructure such as roads, railways, airports | Direct | Gain or Loss |
| Urban development | Any infrastructure built for human settlement (Cities, towns, housing, parks...) | Direct | Loss |
| Mapping error | No observed change as indicated by the model or a transition between two intertidal ecosystems | / | / |
| Unidentifiable | No clear change observed or too much uncertainty in the assessment due to poor-quality images or insufficient imagery | / | / |

We classified each driver type into two overarching categories of change. Direct drivers of tidal wetland extent change were those where the change (either loss or gain) was directly attributable to an event at the location of the change (Angelsen et al, 2018 ; Murray et al, 2022). In these cases the cause of tidal wetland loss or gain was directly visible in high resolution imagery following the change event (such as aquaculture or plantings). Conversely, indirect drivers were processes that influence losses or gains of tidal wetlands but were not observable in high resolution imagery at the location of the change. These change events where no direct driver can be observed were assumed to be the result of indirect activities, and often include the effects of natural coastal processes (e.g. sediment deposition), ecosystem processes (e.g. pioneering vegetation) or the influence of storm events (Spalding 2021). In these cases, the current land cover type occurring at the places of tidal wetland loss or gain were useful for understanding the suite of indirect drivers of change that may be operating at a specific location (Wang et al, 2023).

### 2.2 Data

We used a recently published dataset of Global Tidal Wetland Change (GTWC; Version 1.0) as the basis of this analysis (Murray et al 2022). GTWC data were derived from a global analysis of more than 1 million Landsat 5, 7 and 8 images designed to detect changes (both losses and gains) of tidal wetlands over the period 1999 to 2019. In the GTWC data, loss events were defined as the conversion of tidal wetland ecosystem types (saltmarsh, mangrove and tidal flat) to a non-intertidal land cover type over the period of the analysis. Gain events were defined as the establishment of at least one of the tidal wetland ecosystem types in an area where their presence was not detected in 1999 (Murray et al, 2022). Pixels (30m) representing tidal wetland losses and gains are represented as binary occurrences in data bands ‘loss’ and ‘gain’. To investigate the relative impact of drivers per ecosystem type we used the data bands “lossType” and “gainType” that depict the type of tidal wetland ecosystem lost or gained in each change event (tidal flats, mangrove, or tidal marsh) (Figure 1).

**Figure 1:**
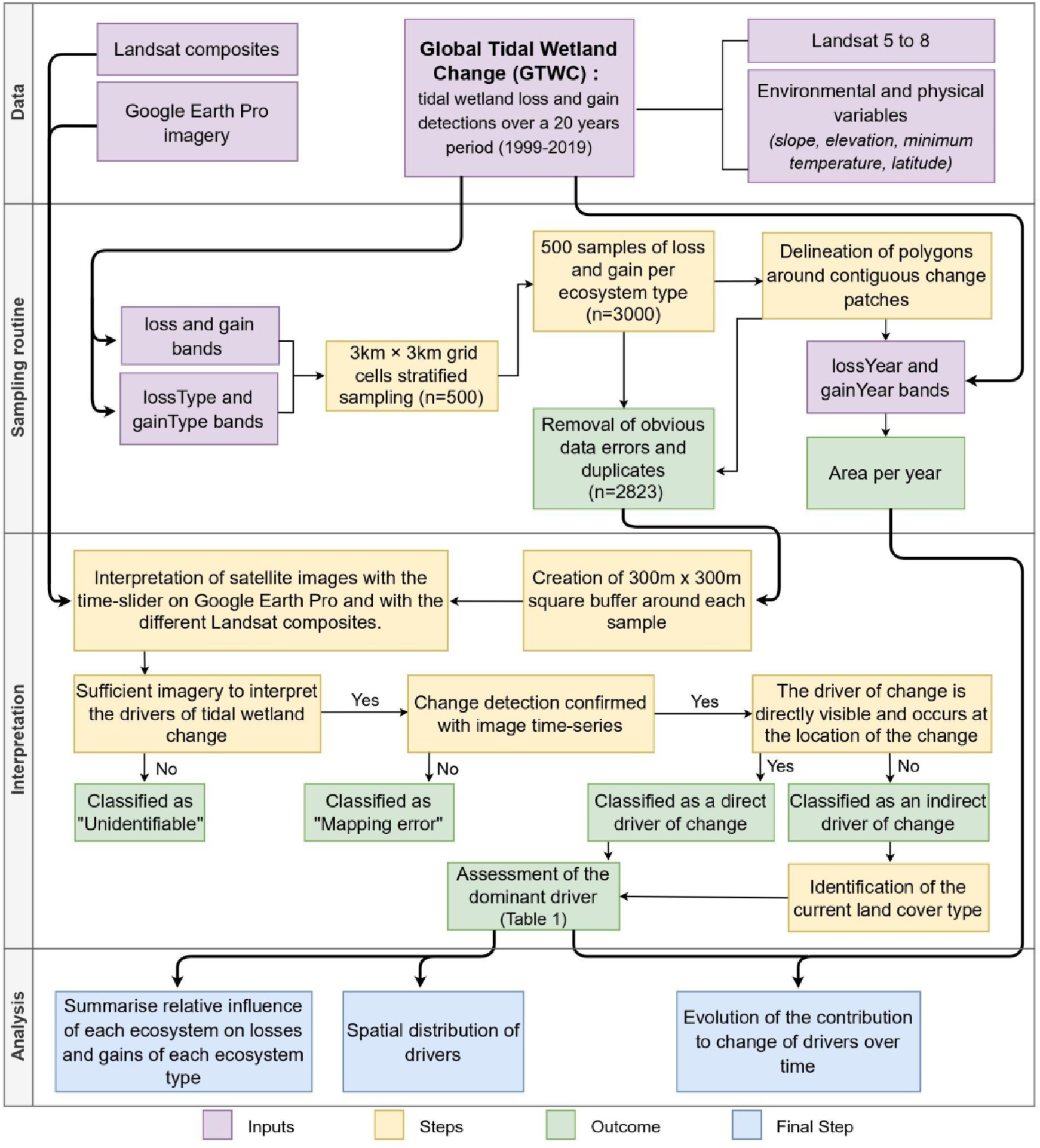
Analysis pipeline for estimating the dominant drivers of global tidal wetland change.

To estimate drivers of change of tidal wetlands across the full extent of the GTWC data 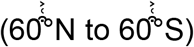, we developed a globally distributed, stratified random sample of detected losses and gain events (Figure S1). We used the same global sampling method developed by Murray et al (2022) but expanded the number of change events sampled. Specifically, we randomly selected 3 km×3 km grid cells (n=500) placed over the GTWC data, using a weighted probability sampling approach that was proportional to the area of tidal wetland change estimated in each grid cell.

Within each selected grid cell, we randomly sampled loss or gain samples of each ecosystem for further analysis, with the number of samples per grid cell informed by the area of each tidal wetland loss or gain per ecosystem. Our sampling approach, which respected the mapping unit of 10 eight-way connected 30m × 30m pixels from the GTWC, yielded 500 samples of tidal wetland loss and 500 samples of tidal wetland gain per ecosystem type, totaling 3000 globally distributed samples for the driver interpretation. A post-sample review removed 8 samples from the sample set due to data errors. Each sample included information propagated from the GTWC data, such as the type of change event (loss or gain), the ecosystem type (tidal marsh, mangrove, or tidal flat).

To explore the temporal contribution of each driver to change and estimate the rate at which different tidal wetland change drivers operate, we used the data bands ‘lossYear’ and ‘gainYear’ from the GTWC. These bands represent per-pixel the year that a loss or gain event was first detected by the analysis. To estimate the rates of operation of each driver, we created polygon features representing the boundary of the loss or gain zone surrounding each sample on the GTWC. For that, adjacent pixels to each sample were grouped to form the polygon. 169 duplicates (polygons with 2 or more samples within their boundaries) were removed. Thus, our final sampled dataset for further analysis consisted of 2823 samples. We then computed the area of change per time period (2002–2004, 2005–2007, 2008–2010, 2011–2013, 2014–2016, 2017–2019) for each polygon.

### 2.2 Driver interpretation

We developed an expert image interpretation procedure to estimate the cause of tidal wetland change for each driver sample in our sample set. For each sample of tidal wetland loss or gain, we used Google Earth Pro (version 7.3.6.9345) to attribute the main driver that contributed to the change. We used the high-resolution imagery available in Google Earth Pro and the historical time slider to view change dynamics before, during, and after each change event to support the expert interpretation (Figure S2).

Four interpreters were tasked with allocating each sample to one of the 18 driver categories in the classification scheme (Table 1). The interpreters used the time-series imagery, as well as features proximate to the pixel of change and features surrounding the pixel of change (as required) to determine their annotations. We recorded driver annotations as the majority driver, which was the driver deemed by the interpreter to be primarily responsible for the change event (Curtis et al, 2018).

In tidal wetland ecosystems, drivers of change operate at a range of spatial scales and often at significant distances from where the initial driver operated. For instance, the ‘sand engine’ in the Netherlands is a sand deposit designed to replenish coastal ecosystems immediately south of the engine itself (Luijendijk et al, 2017). To enable interpretation of this important attribute of tidal wetland change dynamics, driver assessments were conducted at the scale of the 300m x 300m square because we estimated that the single-pixel scale was insufficient to capture the real reasons for a change, possibly underestimating the direct drivers (Figure 2). In 20 years, tidal wetlands may have undergone a succession of several drivers. In these rare cases, the most recent has been taken into account.

**Figure 2:**
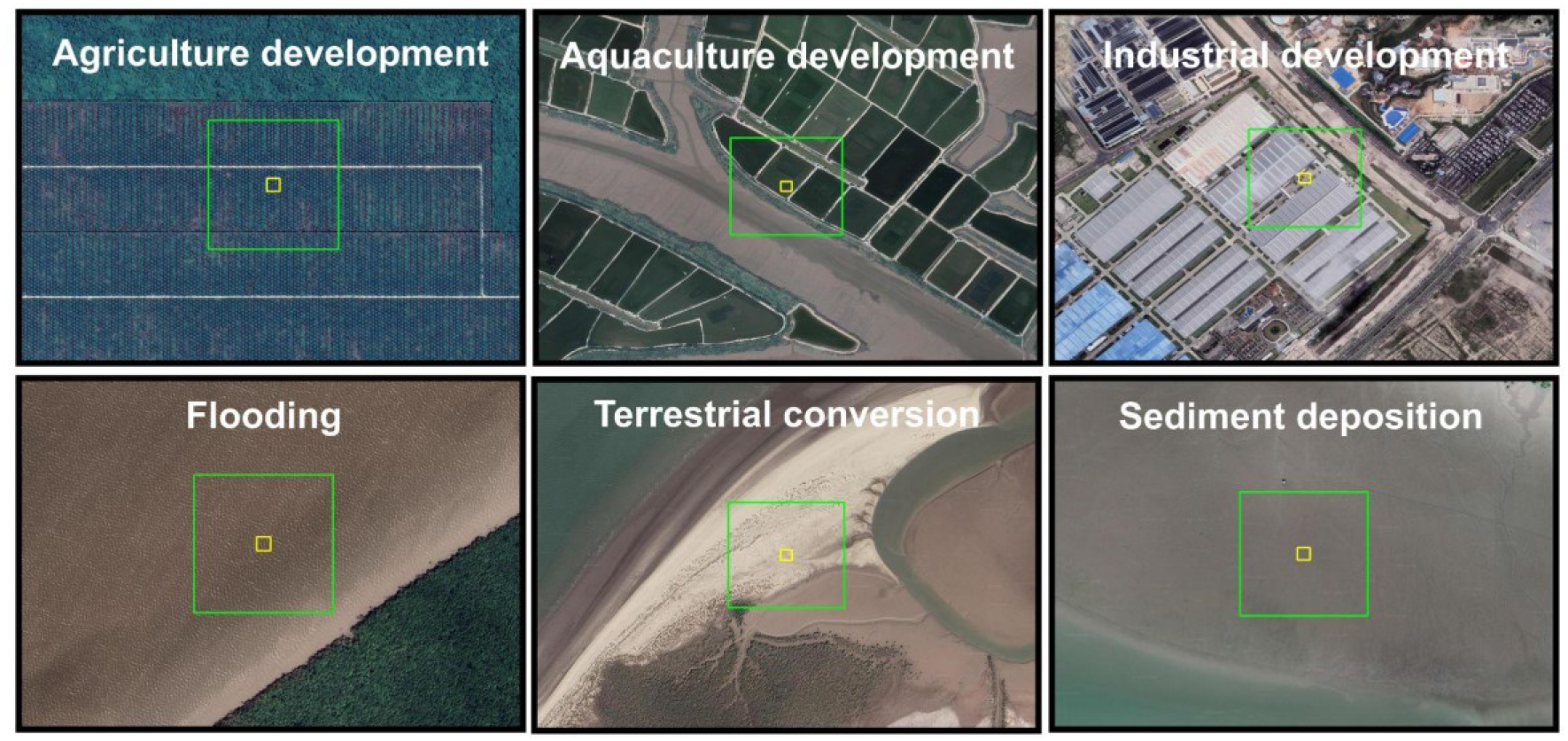
Example of 6 sample pixels and their classifications for samples labelled to a direct driver class (top row) and to an indirect driver class (bottom row) on Google Earth Pro. The yellow square represents the 30m x 30m pixel and the green square represents the 300m x 300m buffer zone used for the assessment.

In many cases, satellite images on Google Earth Pro for the two reference years of the GTWC, 1999 and 2019, were either not available or were of poor quality. Thus to aid our interpretations, we developed several cloud-free Landsat image composites using Landsat data acquired in the years 1999 and 2019 over the full global extent of the GTWC data. Specifically, we visualised the following Landsat composites: (i) the near-infrared band, (ii) a true-colour composite, (iii) a false-colour composite (NIR, Red and Green) and (iv) the standard deviation of the Automated Water Extraction Index over the year period (AWEI, Feyisa et al, 2014).

To ensure interpretation reliability and facilitate cross-communication, 50 samples were collectively assessed by all interpreters. The remaining samples were randomly distributed among interpreters. For cases of uncertain driver assignment (n = 176), a collective review was conducted until consensus was reached. Samples with no observed change or when a driver annotation could not be confirmed, were categorised as “unidentifiable” (n = 347). In cases where no change was observed in reference imagery or transitions between wetland types were apparent, samples were labelled as “mapping error” (n = 246) and excluded from further analysis.

### 2.3 Analysis

We developed analyses to investigate the global distribution and relative influence of drivers on each tidal wetland ecosystem type. To summarise the relative influence of each driver type on global tidal wetland change we calculated the number of change samples by each driver and used Sankey plots to summarise them by ecosystem type. To explore the global dynamics of tidal wetland change drivers, we also conducted a spatial analysis by estimating the proportion of occurrences of each driver on the total loss or gain recorded at the continental scale.

To investigate temporal dynamics of tidal wetland change, we compared the evolution of drivers over time. For that, we calculated the contribution of each driver for each of the time periods. We decided not to represent drivers whose participation in changes has never exceeded 5% in 20 years.

All analyses were conducted in Google Earth Engine (Gorelick et al, 2017), QGIS (version 3.28.3) and R (version 4.2.2; R Core Team, 2022).

## 3. Results

### 3.1 Indirect drivers of tidal wetland change

Our analysis of the globally distributed tidal wetland change samples indicated that indirect drivers of change accounted for a larger proportion of tidal wetland change events (losses and gains combined; 70.9%) than direct drivers (29.1%; Figure 3). Nearly two-thirds of all loss samples (60.7%), were attributed to indirect driver types, accounting for 44.4% of mangrove loss samples, 59.3% of tidal flat samples and 84.3% of tidal marsh samples.

**Figure 3:**
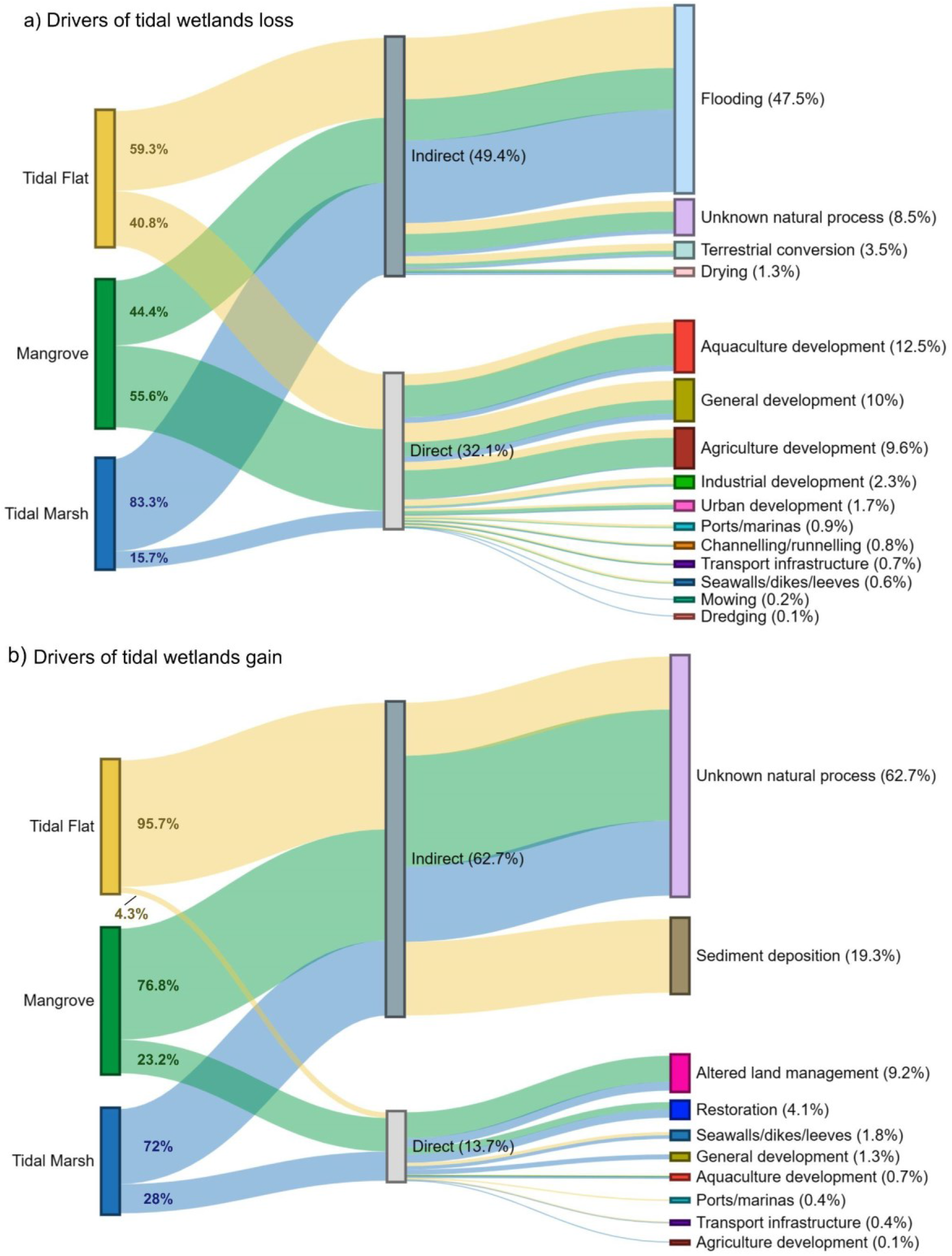
Distribution of drivers of losses (a) and gains (b) according to the type of ecosystem. The coloured percentages represent the distribution of driver types (indirect and direct) for each corresponding ecosystem

Flooding was the most commonly annotated indirect driver of loss across the tidal wetlands combined (47.5%) and for each of the three component ecosystem types (Figure 3(a)). Unknown natural processes that resulted in tidal wetland loss were considered the 5th driver of losses and have been observed on all continents and across all ecosystems.

The global distribution of change samples attributed to indirect drivers was concentrated in North America and Oceania, where more than 90% of losses were due to indirect drivers (94.7% and 92.9%, respectively Figure 4). Conversely, Asia is the only continent where losses were not predominantly attributed to indirect drivers (30.9%). Flooding was the most common annotation of an indirect driver globally, though its occurrence ranged from 23.1% of losses in Asia to 81.6% in North America (Figure 4).

**Figure 4:**
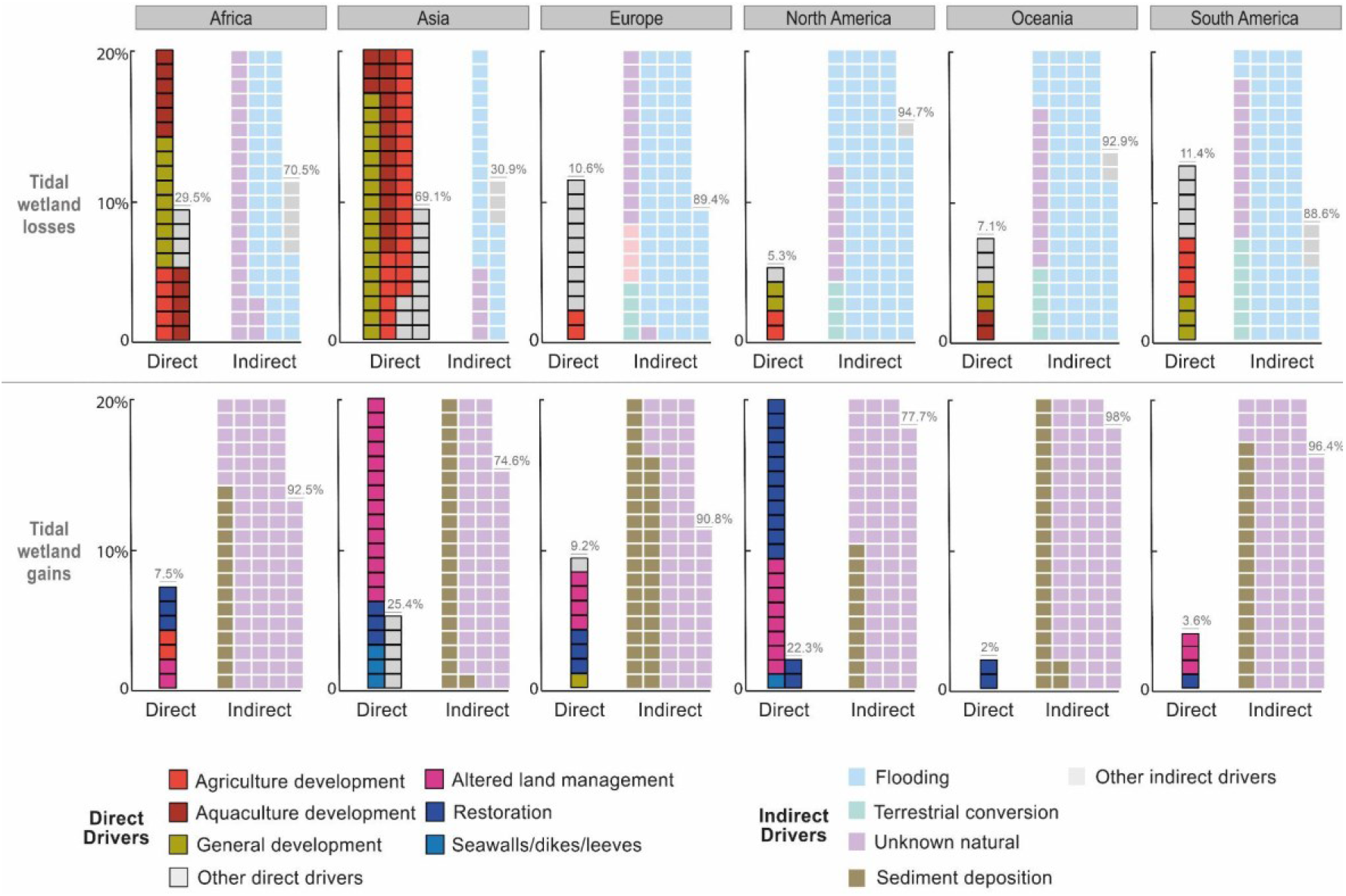
Waffle charts indicating the proportion of driver samples allocated to direct driver (black outline) and indirect driver (no outline). For each continent, the number of cells sums to 100%. Note: Only the 5 most important drivers for each continent have been coloured (independently of driver type) ; the rest of the drivers have been grouped under the category ‘Other drivers’ and represented in grey.

Globally, more than four-fifths (82.1%) of all tidal wetland gains were attributed to indirect drivers of change, with unknown natural processes as the most common annotation (62.7%). Unknown natural processes accounted for three quarters of mangrove gains (76.8%) and 72% of tidal marshes, but only 40.1% of tidal flat gains (Figure 3(b)). Sediment deposition was only attributed to tidal flats, accounting for 19.3% tidal flat gains, ranging from a minimum in North America (9.6%) to a maximum in Europe (35.5%) (Figure 4).

### 3.2 Direct drivers of tidal wetland change

Drivers which were clearly identifiable in satellite imagery at the site of ecosystem change events were identified in 29.1% of the tidal wetland change samples, with 17.9% causing tidal wetland gains and 39.3% of tidal wetland losses. Per ecosystem, direct drivers were identified as the cause of the majority of losses in mangroves (55.6%), but only 40.8% of tidal flats and 15.7% of tidal marshes.

Of the 11 direct drivers identified by our literature review, nine were identified to operate on all of the tidal wetland ecosystem types. Wetland conversion caused by aquaculture (12.5%), general development (10%) and agriculture (9.6%) were the top three direct drivers of tidal wetland loss (Figure 3(a)). These three drivers accounted for 56.8% of Asia’s recorded losses, the continent with the highest number of drivers (n=9) (Figure 4). The two driver types that impacted only one ecosystem type (tidal marshes) were mowing and dredging (0.6% and 0.3% of tidal marsh loss samples, respectively). General development was observed on all continents in the analysis, Agriculture was only absent from Europe and aquaculture only from Oceania.

Direct drivers contributed only a small proportion of tidal wetland gains where no driver accounted for more than 10% of total gains across all ecosystem types. The three most dominant drivers of tidal wetland gain were altered land management (9.2%), restoration activities (4.1%) and seawalls, dikes and levees (2.4%) (Figure 3(b)). Direct drivers contributed the highest proportion (25.4%) to tidal wetland gain in Asia, with altered land management being the most prevalent driver. Restoration is the primary contributor to direct gains in North America, and the sole driver of direct gains in Oceania (Figure 4).”

By ecosystem, direct drivers of gain were identified in 4.3% of tidal flat samples, 23.2% of mangrove samples and 28% of tidal marsh samples. Our analysis also indicated that tidal marshes were subject to greater variability of driver types (eight) than the other two ecosystems, where less than four driver types were identified to contribute to their gains.

### 3.3 Temporal dynamics

Our analysis of the changing prevalence of driver types over time revealed a change in influence of different driver types over time (Figure 5). Flooding was consistently the predominant driver of loss in every time-step of the analysis. Flood-related losses were remarkably high during the first 3 periods of the study (2004 to 2010), constituting nearly half of the total losses in 2004 (49.9%). From 1999 to 2019, losses caused by agriculture and aquaculture decreased from 16.3% to 11.6% and from 12.3% to 5.9%, respectively. (Figure 5). In contrast, the impacts of general development on tidal wetland loss increased from 8.8% of losses in 2004 to 24.8% in 2013.

**Figure 5:**
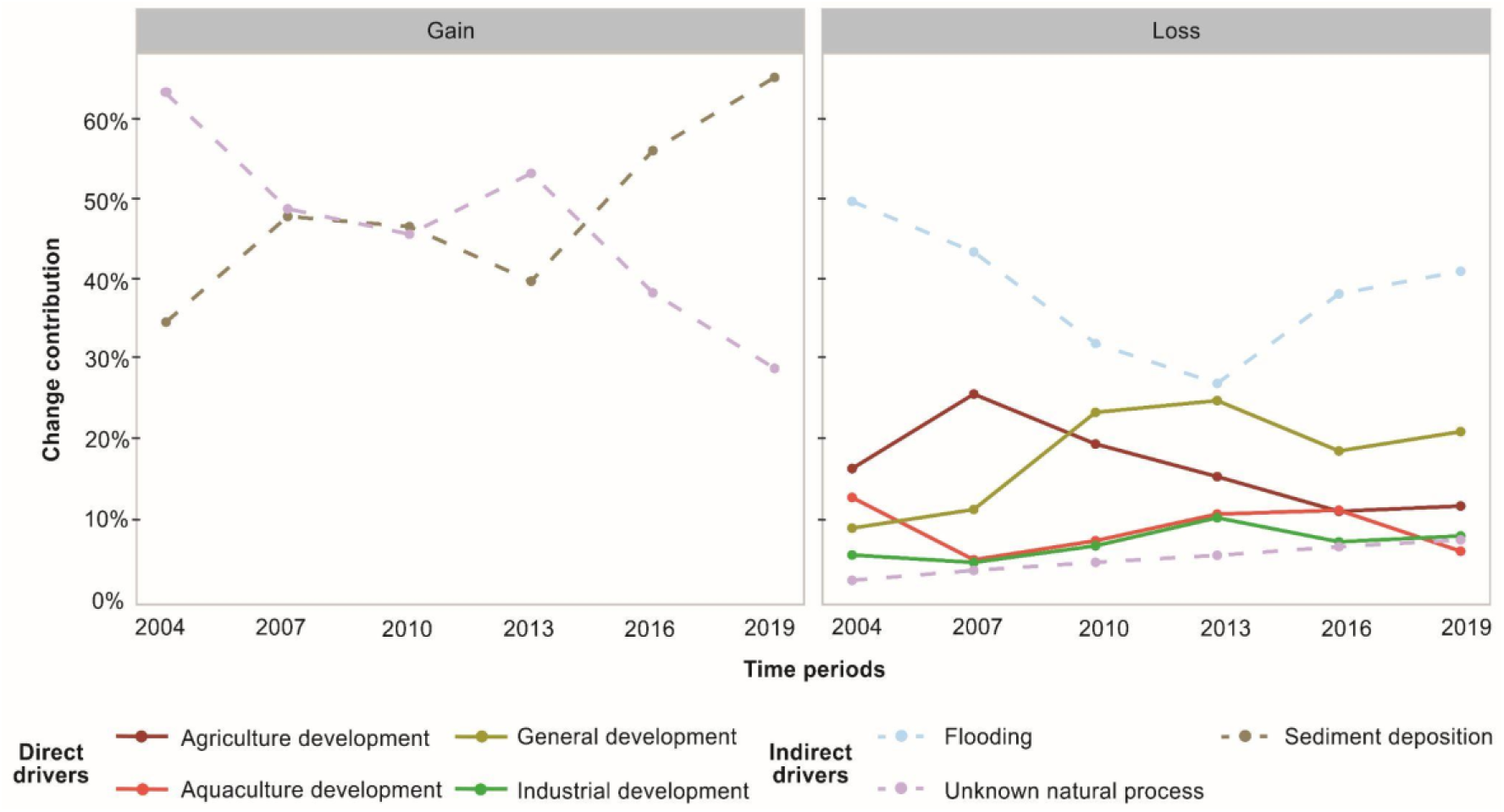
The changing proportion of drivers over time. Note: only drivers that account for more than 5% of the total samples in at least one time period are shown in this plot.

Sediment deposition and unknown natural processes were the only drivers acting on gains for all time-steps over the 20 year period. The contribution to gains by unknown natural processes decreased from 63.3% in 2004 to 28.8% in 2019. Conversely, gains from sediment deposition steadily rose over time, accounting for 65.3% of total gains in 2019 (Figure 5). Throughout our analysis period, direct drivers of gains did not contribute more than 5% of gains in any single time-step. Direct drivers of gain, including general development (maximum of 4.5% of gains in 2013) and restoration activities have remained consistently low over the analysis period (<1% of total gains throughout; Figure 5).

## 4. Discussion

### 4.1 Worldwide driver dynamics of tidal wetland losses

Across all continents and ecosystem types, flooding was the most attributed driver class for losses. The loss of tidal wetlands by transition to permanent water can be the result of various processes including subsidence, compaction, erosion and direct impacts of severe storms (Mazzotti et al. 2009; Silva et al. 2014; Lopes et al. 2021). Similarly, the observable response of tidal wetlands to rising sea level is also conversion to permanent water, a complex process that can be exacerbated by a range of other factors, including an inability to migrate toward higher elevation areas (e.g., beach compression; Gracia et al. 2018; Trenhaile 2018; Lithgow et al. 2019).

The global phenomenon of rising sea levels may already be causing widespread drowning of tidal wetlands. Lin et al. (2024) estimated that 12% of China’s tidal wetlands have transitioned into open water, with erosion as the primary cause. Similarly, Saintelan et al. (2023) found that most in-situ tidal marsh sites monitored by remote sensing models show increased surface water presence. Additionally, they reported difficulties in analysing surface water changes in mangrove ecosystems due to canopy cover confounding satellite observations (Saintelan et al. 2023). Our high-resolution imagery approach, which includes contextual information and time-series analysis, found that the proportion of mangrove samples annotated as flooding to belower than other tidal wetland types. This is likely due to mangroves’ capacity to trap sediment and maintain elevation despite rising sea levels (Mcleod et al. 2011 ; Morris et al, 2023 ; Saintelan et al, 2023).

Aquaculture and agriculture were the primary direct drivers of mangrove and tidal marsh loss and ranked second and third, respectively, for tidal flat loss. These production activities have extensively impacted tidal wetland ecosystems. Aquaculture, such as shrimp and fish farming, has expanded significantly to meet global food demands since 1999, often requiring pond construction in low-elevation, tidally connected areas (Neiland et al. 2001; Goldberg et al. 2020; Ofori et al. al. 2023). The majority of these activities were concentrated in Asia, where over four-fifths of the world’s aquaculture and 90% of rice production occur (De Silva and Davy, 2010 ; Bandumula, 2018 ; FAO, 2020).

We found that between 2004 and 2019, tidal wetland losses attributed to general development surged almost fourfold. This category encompasses on-going activities like tidal wetland reclamation for urban or industrial purposes that typically occur in proximity to existing cities and highly populated coastal areas (Sengupta et al. 2023). From 1980 to 2017, around 1250 km^2^ of coastal land was claimed by 16 megacities (Sengupta et al. 2018). Furthermore, land reclamation and high population density are frequently associated with an increase in detrimental anthropogenic impacts like pollution on the surrounding ecosystems (Alongi 2002; Turschwell et al. 2021).

### 4.2 Worldwide driver dynamics of tidal wetland gains

Unlike the drivers of loss, the majority of drivers of global tidal wetland gains were indirect, highlighting the natural dynamic of these ecosystems to establish in areas they didn’t formerly occur. Mangroves and tidal marsh species exhibit a high colonisation capacity due to the persistence of their propagules (Marchand 2003). Sediment supply, from rivers or human activities, is crucial for offsetting sea level rise by supporting wetland expansion and elevation maintenance, (Kirwan et al., 2011, Goslin et al, 2022). Our findings reveal an increasing contribution of sedimentation to gains, likely influenced by climate change-induced factors such as heightened and prolonged rainfall patterns (Beuselinck et al. 2002; Lopes et al. 2021). Adjacent anthropogenic factors like deforestation can also contribute to sediment availability (Crain et al. 2009; Syvitski et al. 2009).

Direct drivers of gain, such as restoration activities and engineered structures that influence sediment deposition, were prevalent across various ecosystem types. Mangroves and tidal marshes saw substantial gains (23.2% and 28% respectively) from these activities, while tidal flats experienced minimal impact (less than 5%). Tidal wetland restoration activities were observed on all continents but predominantly in North America. However, their overall contribution remains low, possibly due to unsuccessful projects, often misaligned with environmental characteristics (Kodikara et al. 2017 ; Worthington and Spalding 2018 ; Lovelock and Brown 2019). Yet, studies suggest that well-managed restoration actions have demonstrated sustained effectiveness over time (Liu et al. 2020; Damastuti et al. 2022).

Despite observing tidal wetland restoration leading to gains, many drivers linked to gains were also tied to losses, including agriculture, aquaculture, and general development. Sediment dynamics strongly influence the locations of tidal wetland gain areas. For instance, gains in all tidal wetlands areas often coincide with the construction of seawalls and other hard structures in reclamation developments. Yang et al. (2008) identified slight increases in available sediments immediately adjacent to a man-made seawall. The establishment of these ecosystems where losses occur underscores the importance of monitoring net tidal wetland changes (Wu et al, 2018).

## Limitations and conclusions

While our analysis enhances the understanding of tidal wetland change drivers, limitations exist due to the absence or poor quality (e.g. high cloud cover) of high-resolution time-series satellite imagery, hindering confirmation of change or driver attribution. Despite advancements over the past decades, publicly available global-scale image databases remain limited.

Identifying a single factor driving tidal wetland change was complex and time-consuming, given its multifaceted nature. While our analysis focused on primary drivers per sample, local scale impacts like reclamation can also interplay with larger-scale processes like sea level rise, and future work should focus on disentangling these drivers. Remote sensing methods lack the resolution to extricate drivers, leading to relatively simple classification schemas (less than 5 classes) (Richards and Friess, 2016 ; Curtis et al, 2018 ; Goldberg et al, 2020). Although our interpretation approach enabled the identification of more subtle drivers, factors like invasive species (e.g., *Spartina alterniflora*), couldn’t be taken into account despite their known influence (Liu et al., 2018). Field-based studies remain crucial for understanding global tidal wetland change, with future efforts aimed at synthesising decades of research findings.

Our study successfully identified 18 drivers of tidal wetland change, offering valuable insight into the relative impact of a wide range of factors, including natural processes, climate change effects, and human pressures, spanning a two-decade period. This study also shows that it would be necessary, in order to be more consistent, to use a standardised taxonomy of drivers of coastal change dynamics as proposed by Lucas et al, 2022. However, it also underscores the persistent need for refining detailed, accurate, and precise spatial models of tidal wetland change. These models should enhance the frequency of change observations while accommodating the inherent dynamic nature of tidal wetlands. Developing a precise understanding of the threats and mechanisms influencing the distribution of these declining ecosystems, is imperative for implementing effective conservation and restoration measures into the future.

## Supplementary Information

**Figure S1:**
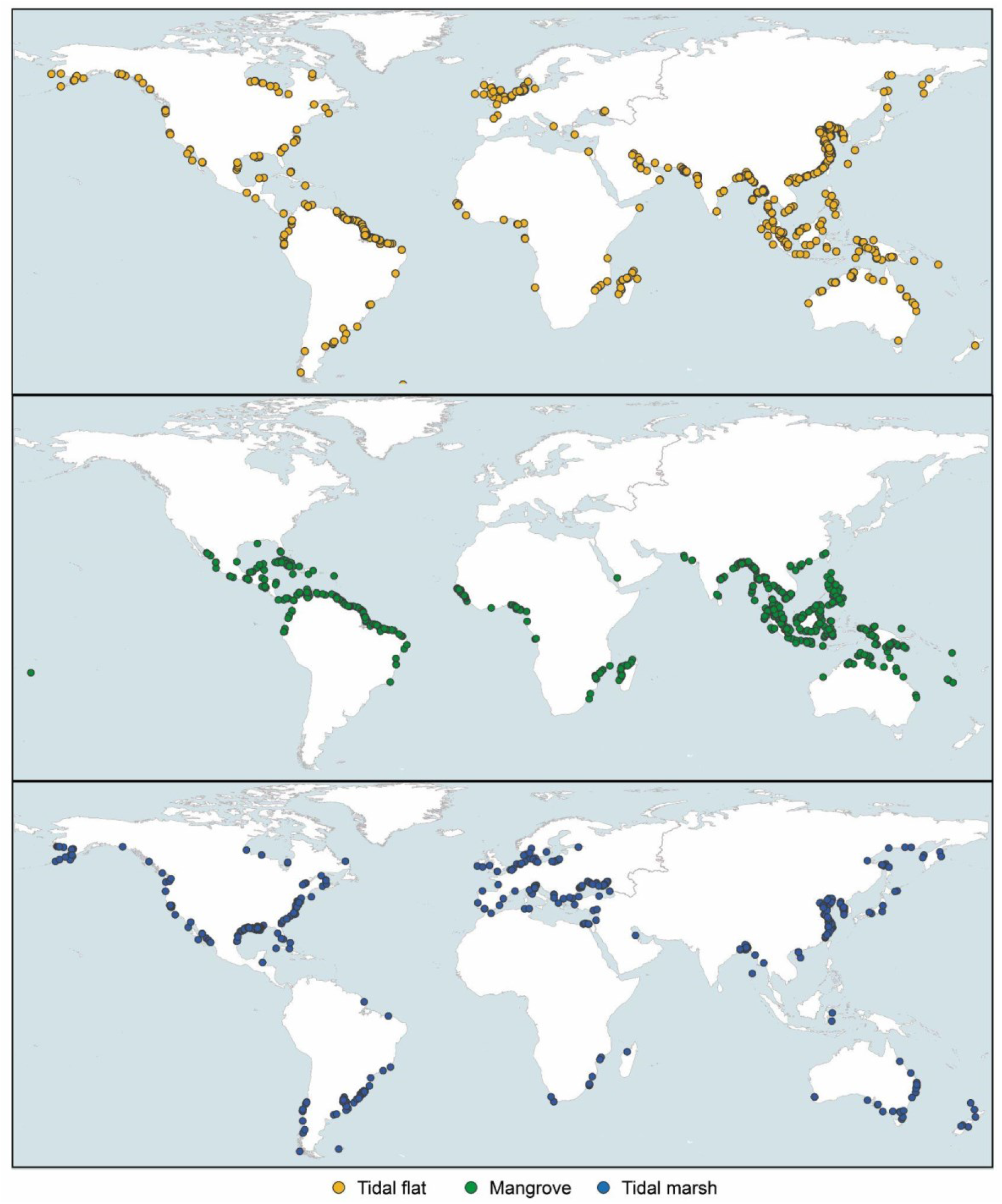
The global distribution of the gains and losses of tidal wetlands dataset (n = 2992) which was randomly sampled from the Global Intertidal Change map (Murray et al. 2022)

**Figure S2:**
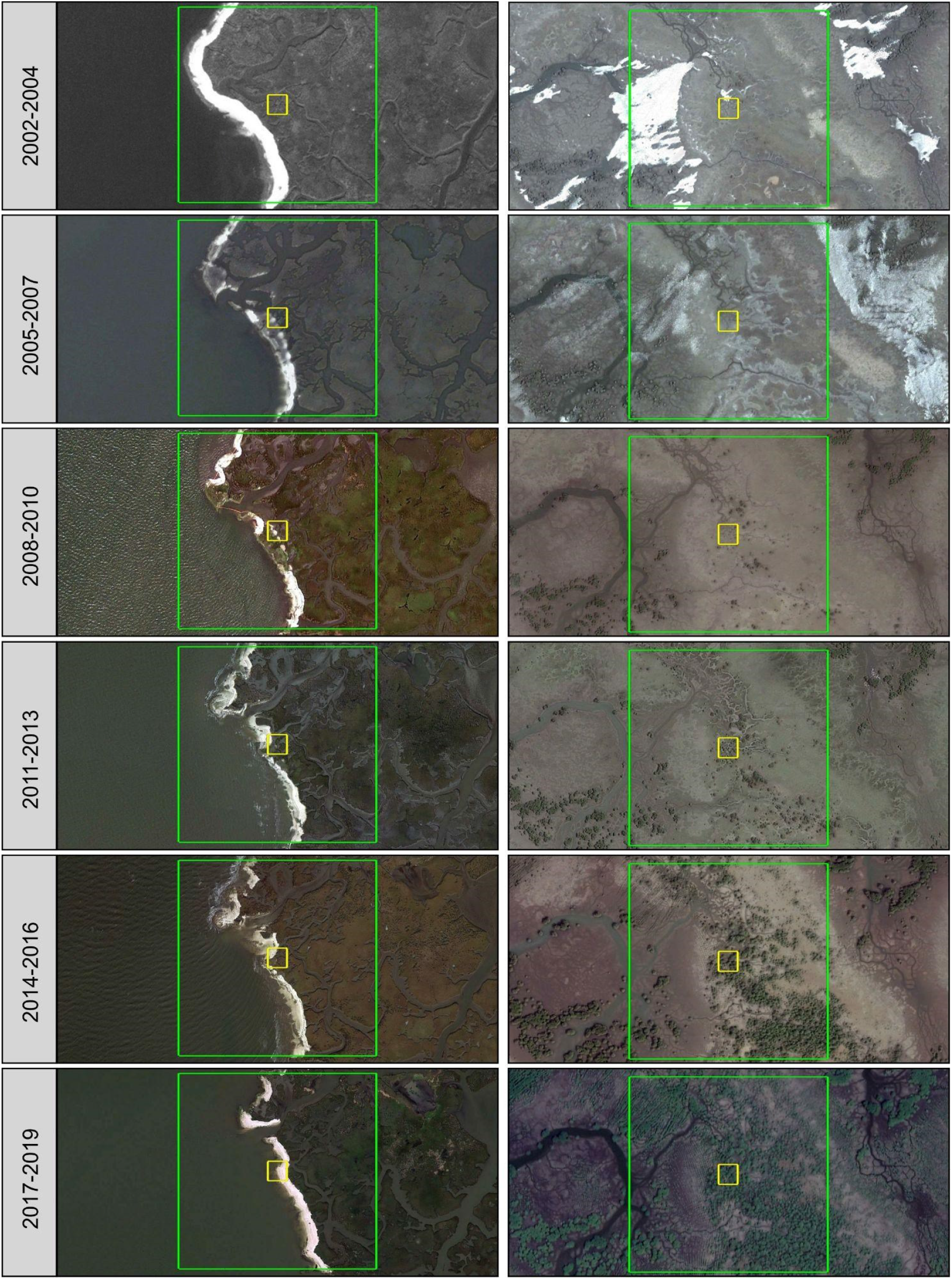
Visualisation of the temporal evolution of land cover type on Google Earth Pro and identification of the driver behind the observed change for one loss sample classified as flooding (left column) and one gains sample classified as restoration (right column). The yellow square represents the 30mx30m pixel and the green square represents the 300mx300m buffer zone used for the assessment.

